# Integrative Multi-omics Deciphers the Assembly Rules of the Human Skin Microbiome and Highlights Lipid-Mediated Niche Partitioning

**DOI:** 10.1101/2025.09.04.674196

**Authors:** Huizhen Chen, Wei Luo, Zhiming Guo, Zhiming Li, Jianju Feng, Shensong Cao, Qi Wang, Zhenglong Gu, Jean Krutmann, Jiucun Wang, Lianyi Han, Yangfan Guo, Jingjing Xia

## Abstract

Understanding the ecological assembly mechanisms of microbial communities is critical for deciphering host-microbiota interactions, however, such processes remain poorly characterized in cutaneous ecosystems. By profiling a non-industrialized rural cohort, this study deciphers the assembly rules governing the human skin microbiome and reveals an evolutionarily conserved core microbiota structured by niche selection. Microbial α-diversity peaks when *C. acnes* and *M. osloensis* reach equilibrium, supporting the Intermediate Disturbance Hypothesis. Host lipids, rather than aqueous compounds, emerged as the primary drivers of microbial niche partitioning. We further demonstrate that *C. acnes* and *Malassezia restricta* function as core metabolic engineers, expressing diverse lipases that hydrolyze triglycerides (TGs) into free fatty acids (FFAs), thereby maintaining skin surface lipid homeostasis (∼85% TG + FFA). Notably, virtually all skin surface FFAs are microbially derived and function as “public goods”, enabling cross-feeding that supports auxotrophic members and fosters mutualistic interactions. Importantly, *in vitro* coculture experiments confirm that *C. acnes* establish a survival niche for *M. restricta*. Our findings support a paradigm shift from a taxonomic to a functional understanding of the skin ecosystem, underscoring microbial lipid metabolism as a central orchestrator of ecological stability and a promising target for therapeutic intervention.

GRAPHIC ABSTRACT

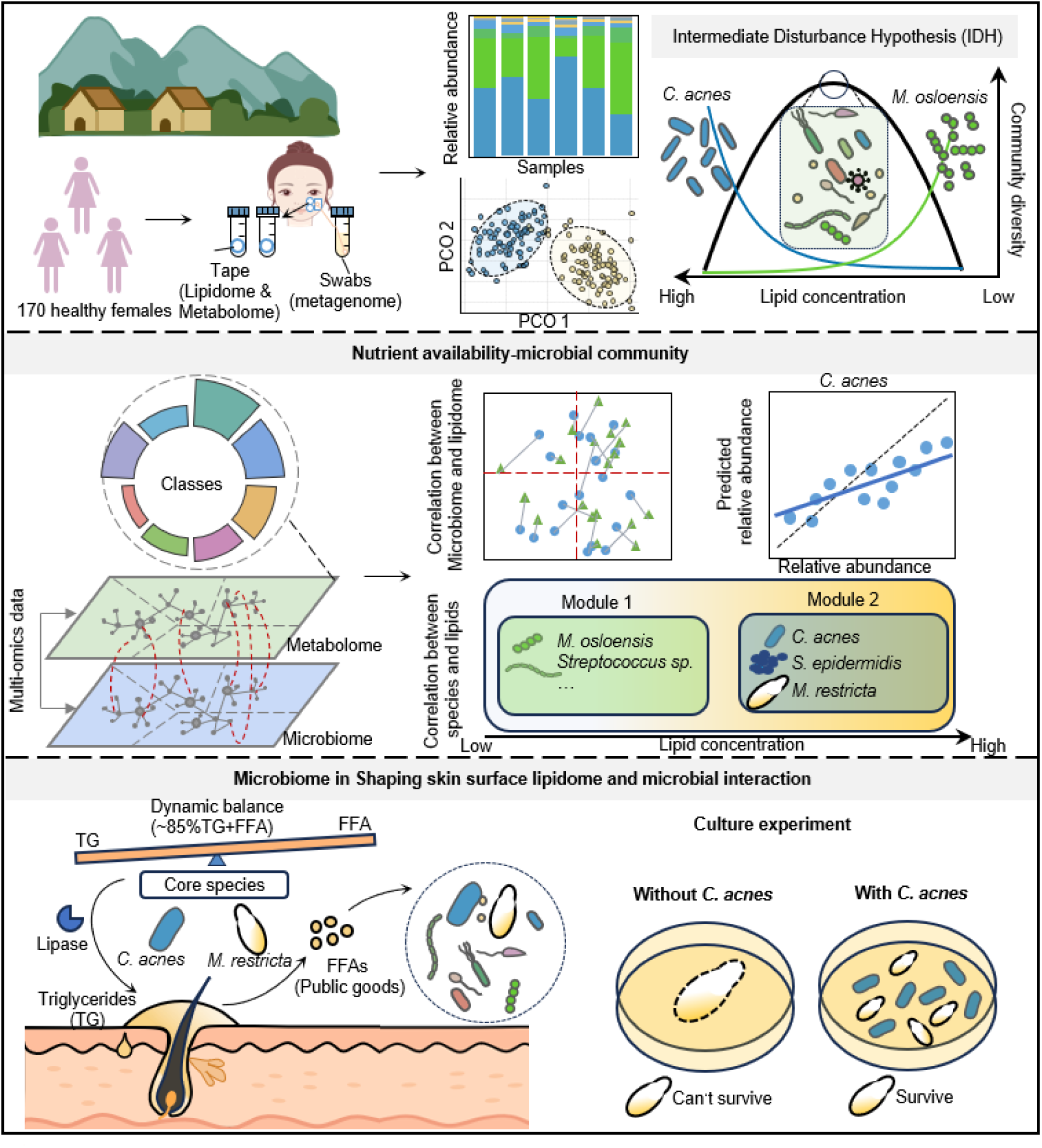

## INTRODUCTION

The human skin harbors a diverse consortium of microorganisms that interact dynamically with the host, forming a complex ecosystem critical to health and disease (*1, 2*). While high-throughput sequencing has enabled comprehensive microbial profiling, fundamental ecological questions, such as those governing community assembly, microbial persistence, and ecosystem stability (*3, 4*), remain largely unresolved. Although metagenomic studies across varied skin sites (e.g., dry, moist, and oily habitats) have offered valuable insights into the role of microenvironments in shaping microbial composition (*5–7*), the absence of systematic and finely resolved analyses has impeded a mechanistic understanding of how local microenvironments drive community assembly. This knowledge gap currently constrains the formulation of predictive models and the development of actionable theoretical frameworks for ecosystem manipulation. Elucidating these mechanisms is critical not only for deciphering the dynamics of skin ecosystems but also for developing targeted treatments for dysbiosis-related conditions such as acne and atopic dermatitis (*8*). Nonetheless, the intricate network of the skin microbiome and its significant interindividual variability pose considerable challenges. A key advance in deciphering this complexity came with the recognition of cutaneous microbial types, termed “cutotypes”, which exhibit distinct network structures, functional profiles, and host associations (*9–11*). This stratification parallels the role of macroscopically discernible “soil types” in terrestrial ecology (*12*). As illustrated in Figure 1a, both skin and soil feature stratified microenvironments, with hair follicles and soil pores, that support dynamic nutrient cycles and habitat heterogeneity. Current theory posits two community assembly processes, deterministic (niche-based) and stochastic (neutral) processes (*13–15*). Deterministic factors include physicochemical conditions such as pH, moisture, and nutrient availability, which filter colonizing taxa, and biotic interactions such as competition, cooperation, and phage predation (*8, 16*). In contrast, stochastic processes reflect random demographic events and dispersal (*17–20*). Although both forces operate (*21*), mounting evidence supports niche-based assembly as the dominant mode on skin (*22, 23*), underpinning its remarkable temporal stability (*24*).

**Figure 1.**
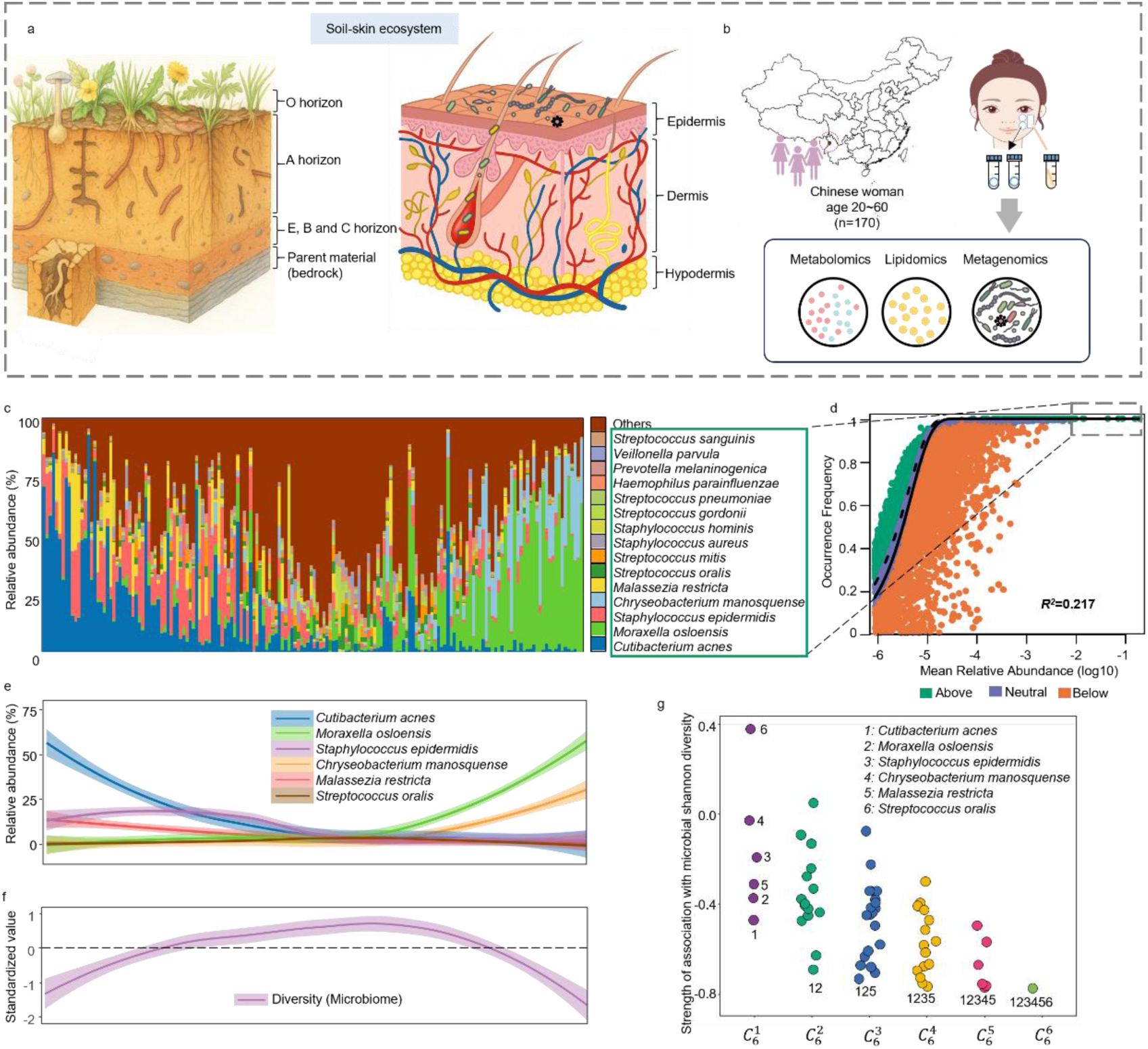
Overview of the facial skin microbiota. a Structural analogy between soil and skin ecosystems. b Schematic of study design. c Composition of top 15 most abundant microbial species. d Sloan neutral model prediction of microbial community. OTUs are represented by data point and colored according to whether the taxon fitted above (green), within (purple), or below (orange) the 95% confidence interval (dotted lines). R2 values (measurement of fit to neutral assembly process). e Fitted Locally Weighted Scatterplot Smoothing (LOWESS) curves with 95% confidence intervals (shaded area) for the abundance of top 6 dominant species. f Fitted Locally Weighted Scatterplot Smoothing (LOWESS) curves with 95% confidence intervals (shaded area) for microbial diversity. g The correlation between the combination of the top six most abundant species and community diversity.

Recent technological advances in metabolomics and the establishment of non-invasive sampling methods (*25*) now enable detailed mapping of the skin’s chemical landscape, the ecological microenvironment in which microbial communities reside and thrive. To decode the interplay between abiotic and biotic factors in this system, we conducted an integrated study of 170 healthy individuals combining shotgun metagenomic sequencing with tape-stripping metabolomic profiling of skin surface compounds (Figure 1b). This multi-omics approach achieves previously unattainable precision in characterizing the niche metabolic microenvironment and microbial interactions. To minimize exogenous confounders, we focused on a cohort from a remote non-industrialized rural area in Yunnan Province, China, where participants have minimal exposure to antibiotics, cosmetics, and environmental pollutants (*26*). This setting provides a unique opportunity to study an “undisturbed” microbial community, likely reflects the intrinsic host-microbe interaction patterns shaped by long-term co-evolution. Through this multi-omics framework, our study aims to unravel the ecological architecture of the skin microbiome, revealing fundamental principles of microbial coexistence and resilience, and offering novel insights for microbiome-based therapeutics.

## RESULTS

### The Skin Microbiome of a Healthy Rural Population

We first characterized the skin microbiota composition in this rural cohort, which exhibited high inter-individual variation (Figure 1c). Core taxa, including *Cutibacterium acnes*, *Moraxella osloensis*, *Staphylococcus epidermidis*, *Chryseobacterium manosquense*, *Malassezia restricta*, and *Streptococcus oralis*, largely overlapped with those in industrialized Shanghai cohort (*9*), suggesting a conserved core structure within the human skin ecosystem. The species *C. manosquense*, however, has thus far been described in only one prior study, which detailed its isolation from a skin swab (*27*).

The Sloan neutral community model (NCM) was applied to evaluate whether skin microbial assembly follows a neutral process (11). The model showed a poor fit to the data (R² = 0.207), with high-abundance taxa such as *C. acnes*, *M. osloensis*, *S. epidermidis*, *C. manosquense*, *M. restricta*, and *S. oralis* consistently fell outside neutral expectations, indicating that their colonization is influenced by niche-specific selection rather than stochasticity. Principal coordinates analysis (PCoA) based on Bray–Curtis distances revealed that the first two axes (PC1 and PC2) explained 20% and 16% of the total variance, respectively (Figure S1a). When samples were ordered along PC1 and microbial compositions were visualized as stacked bar plots, we observed a gradual decrease in the relative abundance of *C. acnes* and *M. restricta* from left to right, while *M. osloensis* and *C. manosquense* showed an opposite increasing trend. *S. epidermidis* displayed a unimodal pattern, first increasing then decreasing, whereas *S. oralis* remained relatively stable without a clear directional trend (Figure 1e). Samples at the two extremes of PC1 were characterized by distinct species enrichment: one end dominated by *C. acnes*, and the other by *M. osloensis* (Figure 1c), consistent with our previous findings on the two distinct skin microbial cutotypes identified in the healthy Shanghai cohort (*9*).

Microbial alpha diversity (Shannon index) along PC1 exhibited a unimodal distribution, peaking when the relative abundance ratio of *C. acnes* to *M. osloensis* approached 1:1, suggesting maximal community diversity under intermediate dominance conditions (Figure 1f). This pattern aligns with the core prediction of the Intermediate Disturbance Hypothesis (IDH), implying a dynamic equilibrium between resource competition and niche partitioning in the skin microecosystem (*28*). Specifically, when either *C. acnes* or *M. osloensis* dominates, competitive exclusion may suppress other taxa, reducing diversity; in contrast, transitional states may create niche opportunities that facilitate coexistence (*28*).

To further quantify the contributions of specific species to community diversity, we conducted correlation analyses between all possible combinations of the top six most abundant taxa (63 combinations in total) and alpha-diversity indices. All combinations that included the pair of *C. acnes* and *M. osloensis* showed significant negative correlations with alpha diversity (Figure 1g). In particular, the *C. acnes*–*M. osloensis* pair exhibited the strongest explanatory power (R² = 0.692, p = 3.1 × 10⁻²⁵), accounting for 49.8% of the diversity variation. The inclusion of additional taxa only marginally improved model fit. These results strongly indicate that *C. acnes* and *M. osloensis* are key drivers of diversity variation within the community and suggest that a balanced coexistence between these two species may help stabilize the skin microbiome.

### Skin Metabolome Diversity Reveals Lipidome-Driven Assembly of Microbial Communities

Within the framework of niche selection theory, we investigated how substrate availability, as a key ecological driver (*29, 30*), influences microbial community assembly through quantitative profiling of the skin surface metabolome. The skin harbors a variety of low-molecular-weight, water-soluble metabolites, including amino acids, sugars, and carboxylic acids (Figure 2a), as well as electrolytes, elements, and inorganic ions, all of which are essential for microbial growth (*31*). In our cohort, a total of 458 water-soluble compounds were identified through semi-quantitative metabolomics (Figure 2a). The coefficients of variation (CVs), calculated as the ratio of standard deviation to mean across individuals, ranged from 0–30% for these compounds, with the majority exhibiting CVs below 10%. This indicates that the water-soluble chemical environment of the skin is highly conserved across individuals, likely providing a low-variability baseline substrate environment (Figure 2b).

**Figure 2.**
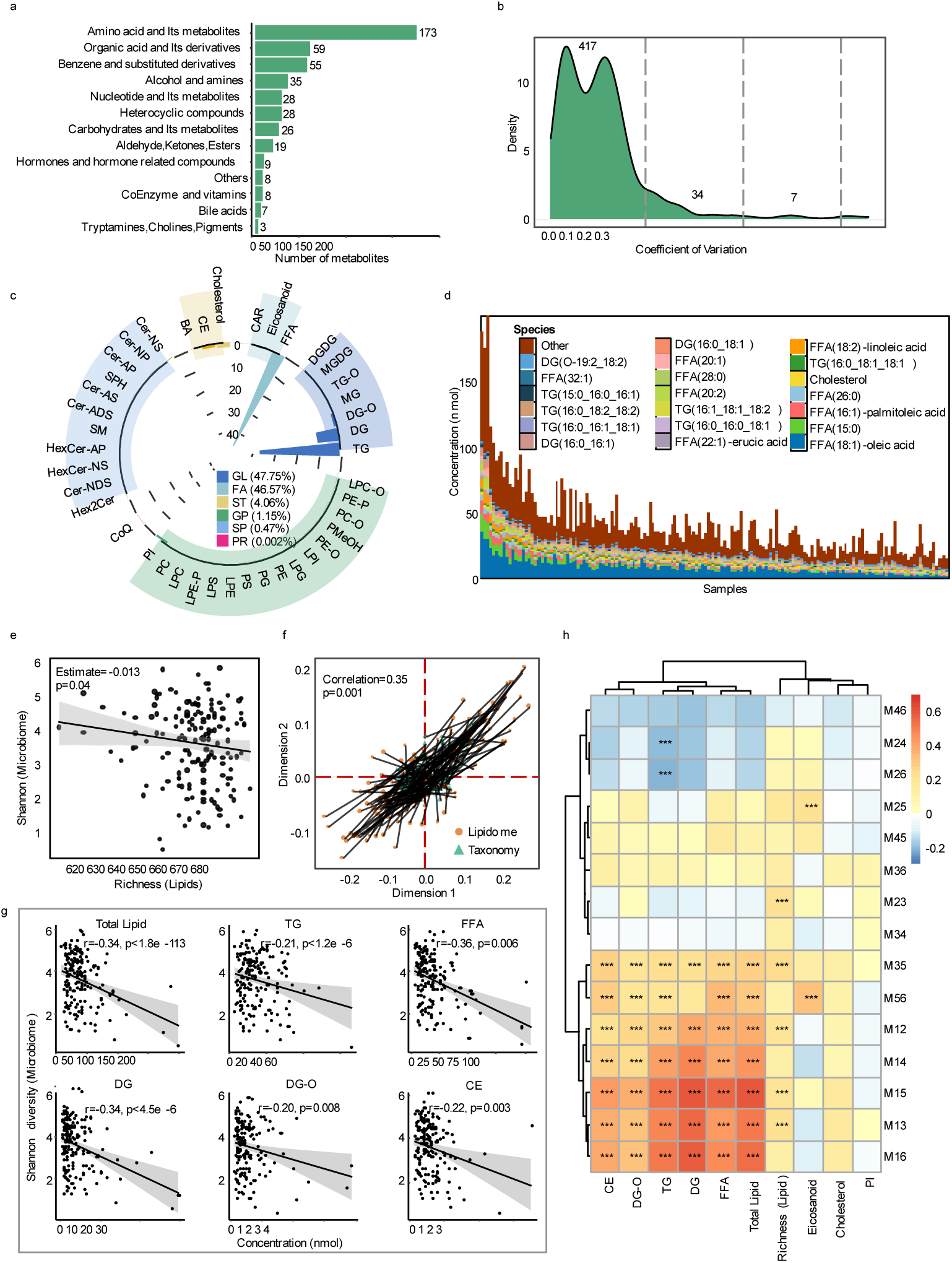
The landscape of the facial skin surface metabolites. a The bar plot represents the counts of each class across all samples. b Distribution of metabolite coefficient of variation (CV). c The circle plot represents the relative abundance of lipid subclasses within each main class and the number depicts the average proportion of each main class in all samples. d Proportion of top 8 lipid classes. e Correlation between the concentration of Lipid Richness and microbial diversity. f Procrustes analysis showing congruence between the microbiome and lipidome. g Spearman’s correlation between the concentration of Top 8 lipid classes and community diversity. h The correlation between the combination of the top six most abundant species and Top 8 lipid classes and lipid richness.

Absolute quantitative lipidomics detected 693 lipid species (Figure S2a). Based on the LIPID MAPS Structure Database (LMSD), these lipids were categorized into six major classes: 421 glycerolipids (GL; 47.75% relative abundance), 48 fatty acids (FA; 46.57%), 36 sterol lipids (ST; 4.06%), 53 glycerophospholipids (GP; 1.15%), 134 sphingolipids (SP; 0.47%), and 1 isoprenoid lipid (PR; 0.002%) (Figure 2c). These were further subdivided into 41 subclasses (Figure 2c), with triglycerides (TG) within GL and free fatty acids (FFA) within FA showing the highest relative abundances. Together, FFA, TG, DG, cholesterol, CE, DG-O, PI, and eicosanoids accounted for 99.0% of total lipid content (Figure S2b-c). In sharp contrast to water-soluble metabolites, the lipidome exhibited substantial inter-individual variability in both total abundance and composition (Figure 2d). Among the 693 lipids, 15 species with relative abundance >1%, including FFA (18:1), FFA (15:0), FFA (16:1), cholesterol, and TG (16:0_18:1_18:1), collectively accounted for ∼50% of total lipid content (Figure 1i), and these high-abundance species also displayed pronounced individual-specific patterns (Figure 2f).

We next integrated ecological data derived from metabolomic and microbiome profiling. A negative correlation was observed between skin surface lipid richness and microbial diversity (Figure 2e), suggesting that increased lipid richness correlates with reduced microbial alpha diversity. This result contradicts the Resource Diversity Hypothesis, which posits that resource heterogeneity promotes species coexistence, and instead points to the presence of inhibitory mechanisms, such as lipids that impair microbial survival or competitive exclusion enhanced by resource diversity(*32*) (*33*). Procrustes analysis revealed a significant association between the overall metabolomic profile and microbiome composition (p < 0.05; Figure 2f). Among the metabolites, 189 lipids were significantly correlated with microbial alpha diversity, with triglycerides (TG) representing the most prevalent class (Figure S2d). Key lipid classes strongly associated with diversity included TG, FFA, DG, and CE (Figure 2i). Furthermore, pairwise correlation analysis between the top six microbial taxon combinations and abundant lipid classes indicated that lipid richness was significantly and positively linked to these microbial pairs. The strongest association was found between lipid richness and the *C. acnes–M. restricta* pair (Figure 2j), implying that lipids may shape community diversity largely through modulating the abundance of *C. acnes* and *M. restricta*.

Notably, our analyses revealed no significant correlations between water-soluble metabolites and microbial diversity (Figure S2d), potentially attributable to their limited inter-individual variation. This observation suggests that the aqueous cutaneous environment represents a conserved chemical baseline, where small hydrophilic metabolites function primarily as fundamental nutritional resources for microbial communities. In contrast, the highly personalized lipid profile appears to serve as a primary driver of ecological niche specialization, shaping microbe-host interactions through selective colonization pressures. This inference aligns with existing literature highlighting the pivotal role of lipid molecules in host–microbe interactions (*34*) (*35*).

### Metabolomic-Microbial Correlation Analysis Reveals Taxa-Specific Nutritional Niches in Skin Microbiota

To decode lipid-driven microbial selection at finer resolution, we quantified associations between dominant microbial species (top 15) and the skin lipidome. Hierarchical clustering revealed two distinct modules of lipid–microbe associations (Figure 3a). The left module included *C. acnes*, *S. epidermidis*, *M. restricta*, and *S. aureus*, which were positively correlated with a wide range of lipids. The right module comprised *Moraxella osloensis*, *Chryseobacterium manosquense*, *Streptococcus gordonii*, *Prevotella melaninogenica*, *Veillonella parvula*, *Streptococcus oralis*, *Streptococcus sanguinis*, *Staphylococcus hominis*, *Haemophilus parainfluenzae*, *Streptococcus mitis*, and *Streptococcus pneumoniae*, which showed negative correlations with many lipid species. Interestingly, this data-driven modular segregation reflects established microbial trophic strategies. The left module, enriched in canonical lipophiles (*C. acnes* and *M. restricta*), comprises species with well-documented lipid dependence (*34, 36, 37*). Conversely, the right module harbors *Streptococcus* species (*10, 38*) and the oligotroph *M. osloensis* (*39–42*), both adapted to low-lipid niches. This systematic partitioning suggests that observed lipid–microbe correlations encode fundamental nutritional specialization within the skin ecosystem.

**Figure 3.**
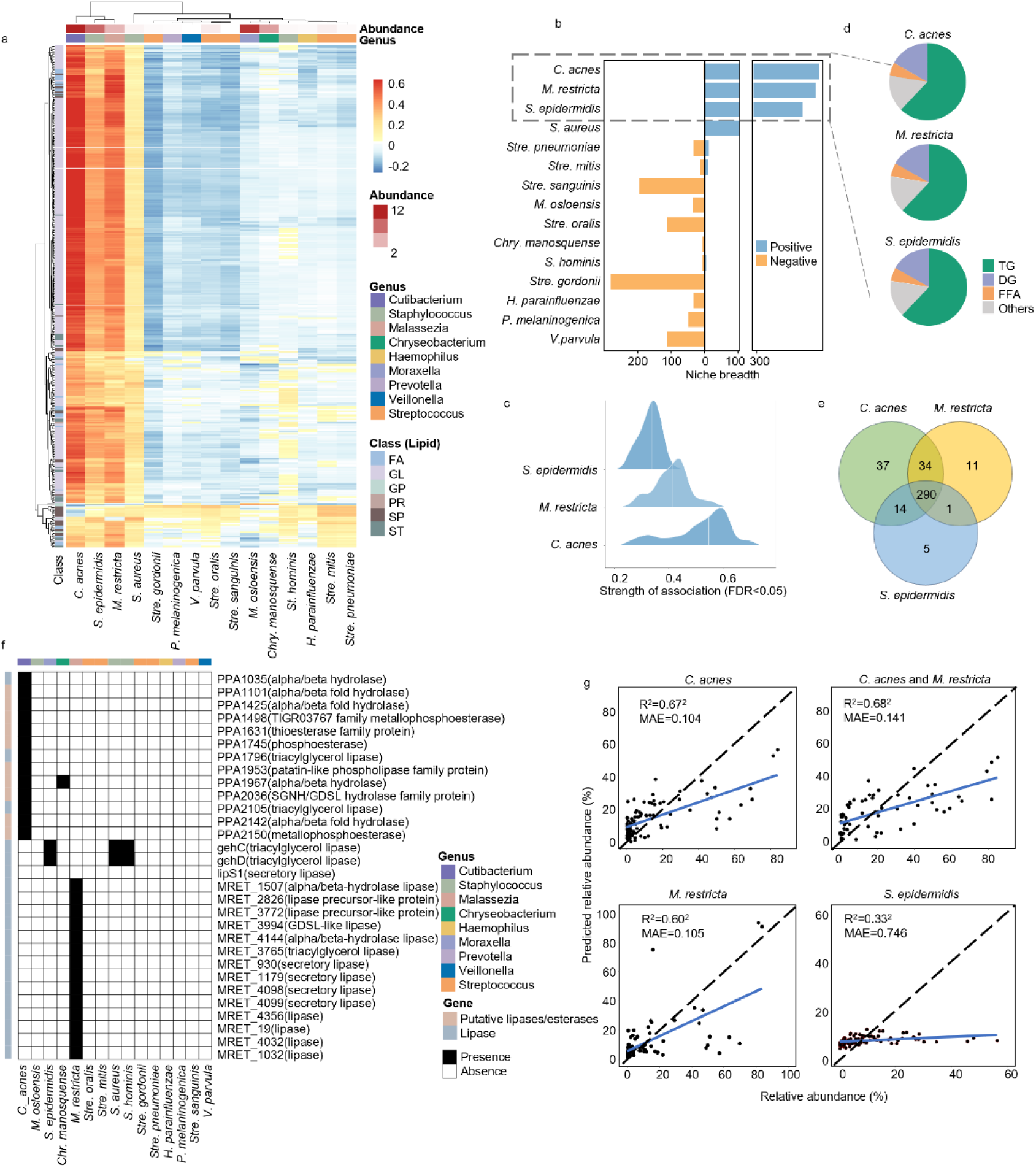
Metabolome–microbiome associations. a Spearman correlation between lipids and species of common genera on the skin. b Distribution of association counts between lipids and species. c Distribution of spearman correlation coefficients between lipid species abundance and lipids. d Main lipid classes associated with the three species. e Venn diagram showing the overlap and specificity of lipid species significantly correlated (p < 0.05) with *C. acnes*, *S. epidermidis* and *M. restricta*. f Presence (black) or absence (white) of gene of lipase and esterase among species. g Predicted relative abundance of *C. acnes*, *M. restricta, and S.epidermidis* and their linear relationship with observed relative abundance. Spearman’s correlation coefficients and mean absolute error (MAE) are denoted.

We subsequently quantified lipid species associations for each microbial taxon as a potential indicator of metabolic niche breadth (*43*). The three predominant lipophilic skin microbes, i.e. *C. acnes*, *M. restricta*, and *S. epidermidis*, demonstrated the most extensive lipid associations (Figure 3b). Notably, *C. acnes* demonstrated both the broadest (Figure 3b) and most unique (Figure 3e) lipid association profile, reflecting its unparalleled metabolic versatility among skin commensals. This adaptive strategy likely underpins its ecological dominance as a keystone lipid-metabolizing species in the cutaneous microenvironment (*34*). This advantage is also mechanistically supported by its extensive repertoire of lipases (Figure 3f) and dominant association with 34 of 37 TG/DG/MG species (Figure 3e), confirming its primary role in skin lipid turnover. While *M. restricta* and *S. epidermidis* showed more limited lipid associations (Figure 3b-c), all three species shared preferential interactions with FFA, TG and DG lipid classes (Figure 3d). Notably, 16 of 29 detected FFAs were commonly associated with these species, suggesting FFA-mediated cross-feeding may represent an important ecological mechanism in the skin microbiome.

To assess the actual lipid-degrading potential of these microbes, we examined the presence of lipase/esterase genes in their genomes using NCBI annotations. Consistent with the observed correlations, both *C. acnes* and *M. restricta* genomes contain multiple lipase/esterase genes. In addition to the previously identified secreted triglyceride lipase gene PPA2105, *C. acnes* harbor several other lipase/esterase genes. The *M. restricta* genome contains 14 annotated lipase genes. *S. epidermidis* possesses fewer lipase genes, while other *Staphylococcus* species such as *S. aureus* and *S. hominis* also contain lipase genes. Although *Chr. manosquense* harbors hydrolase genes, no lipase genes were detected, and the remaining species lacked annotated lipase/esterase genes (Figure 3f). We also reanalyzed previously published raw data obtained from two-dimensional electrophoresis coupled with matrix-assisted laser desorption/ionization mass spectrometry (2-DE-MALDI-MS) (*44, 45*). This data originated from *in vitro* culture supernatants and was used to characterize the secreted proteomes of *C. acnes* and *M. restricta*. *C. acnes* was found to secrete two major lipases: GehA (PPA2105) and GehB (PPA1796) (*44, 46–48*). In *M. restricta*, three secreted lipases—MrLip1, MrLip2, and MrLip3—were purified (*49*). These lipases exhibit distinct substrate specificities and optimal activity conditions (Table S2). Notably, the optimal conditions for GehA and MrLip1 closely resemble those of the healthy skin surface (*50, 51*). MrLip1 shows peak activity at 34°C and pH 5.0 (*52*), while *C. acnes* lipases are stable across acidic pH 5– 8 and have an optimal temperature of 35°C (*51*). GehA can hydrolyze TG, MG, and DG non-specifically (*51*), whereas MrLip1 and MrLip2 are active only against MG and DG (*50*). MrLip3, however, can act on TG, MG, and DG (*50*). Based on these findings, we propose that *C. acnes* can sequentially degrade TG into FFA, while *M. restricta* preferentially hydrolyzes DG and MG to produce FFA.

Given the close relationship between metabolites and microbial abundance, we next developed predictive models using metabolite data to estimate the abundance of *C. acnes*, *S. epidermidis*, and *M. restricta*. *C. acnes* abundance was predicted with the highest accuracy (R² = 0.43, MAE = 0.104; Figure 3g), indicating that skin metabolites explain 43% of its abundance variation. *M. restricta* showed moderate predictability (R² = 0.36, MAE = 0.105), while *S. epidermidis* was poorly predicted (R² = 0.11, MAE = 0.746; Figure 3g). Predicting the combined abundance of *C. acnes* and *M. restricta* did not substantially improve model performance (R² = 0.46, MAE = 0.141; Figure 3g). These results confirm the role of metabolites in shaping microbial community structure and underscore the close ecological relationship between *C. acnes* and the skin lipid microenvironment.

### Role of the microbiome in shaping skin surface lipidomics

Despite the substantial inter-individual variability in skin surface lipids, the combined proportion of triglycerides (TG) and free fatty acids (FFA) remained remarkably stable at approximately 85% across individuals (Figure 4a). These lipid classes exhibited a strong inverse correlation (r = –0.946, p < 6.5 × 10⁻⁸⁴; Figure 4b), suggesting tight metabolic coupling (Figure 4a).

**Figure 4.**
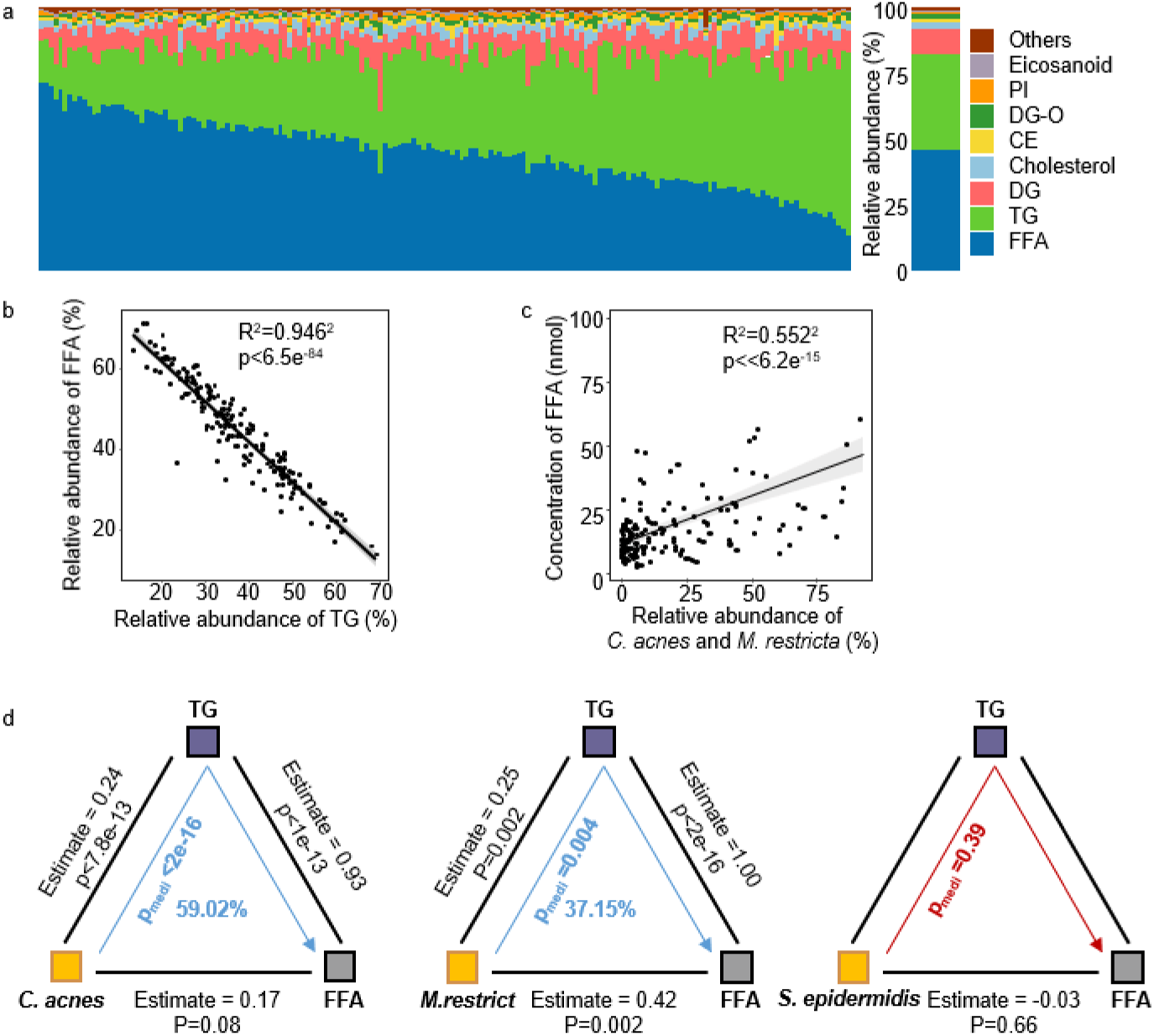
Microbiome in Shaping skin surface lipidomics. a Composition of top 8 most abundant lipid classes. b Correlation of relative abundance between FFA and TG in the population. c Correlation between relative abundance of *C. acnes* and *M. restricta* and concentration of FFA in the population. d The mediation effect of *C. acnes*, *M. restricta* and *S. epidermidis* on FFAs via TGs.

Genomic analysis revealed that both *C. acnes* and *M. restricta* possess multiple lipase genes (Figure 3f). Reductions in the combined abundance of *C. acnes* and *M. restricta* were associated with decreased FFA levels and concomitant TG accumulation (Figure 4c). Among all 63 combinations of the six most abundant species, the total abundance of the *C. acnes* and *M. restricta* pair demonstrated the strongest positive correlation with FFA levels (r = 0.55, p < 6.2 × 10⁻¹⁵; Figure 4d), positioning them as the most capable species for catalyzing the conversion of TG to FFA. Although *Staphylococcus epidermidis* also possesses lipase genes, its relative abundance showed no significant correlation with total FFA levels (p = 0.49; Figure S3a).

Causal mediation analysis further demonstrated that *C. acnes* influence FFA levels predominantly through TG modulation rather than direct effects (Figure 4e). While the direct effect of *C. acnes* on FFA levels was minimal and non-significant (p = 0.08), its indirect effect mediated through TG regulation was highly significant (p < 2×10⁻¹⁶), accounting for 59.02% of the total observed effect (Figure 4e). This mechanistic insight aligns with previous *in vitro* findings showing that selective inhibition of *C. acnes* using demeclocycline (DMCT) led to a ∼50% reduction in surface FFAs (*53*), confirming its pivotal role in generating skin-derived FFAs through TG hydrolysis. *M. restricta* showed a similar but weaker effect (37.15% mediation). In contrast, *S. epidermidis*, also known as a lipophilic species, exhibited neither a direct effect on FFA levels nor an indirect effect via TG metabolism (Figure 4e), which aligns with an *in vitro* finding showing that *S. epidermidis* contributes minimally to FFA production (*54*).

Taken together, these results support the conclusion that in addition to the positive selection of suitable species by the lipid environment, microbial metabolic activity significantly remodels the fine composition of skin surface lipids—particularly the dynamic balance between TGs and FFAs. Like other lipids, FFAs on the skin surface originate either from the stratum corneum or from sebaceous gland secretions. Given that the lipids in the stratum corneum—FFAs, ceramides, and cholesterol homologs— are typically present in a 1:1:1 molar ratio (*55–57*), we estimate that stratum corneum-derived FFAs account for only 0.09% to 5% of the total FFAs on the skin surface (Figure S3a). This suggests that over 95% of surface FFAs are derived from sebaceous lipids. However, freshly secreted sebum contains negligible amounts of FFAs (*58*). Therefore, we infer that the FFAs present on the skin surface are primarily generated via microbial metabolism of sebaceous gland-derived lipids. This is consistent with earlier findings that surface FFAs are largely products of microbial hydrolysis of sebaceous triglycerides (*36, 54, 55*). *C. acnes* and *M. restricta* are the principal microbial executors of TG-to-FFA conversion on the skin surface.

### FFAs: A Key Regulatory Factor in the Skin Microecosystem

FFAs are essential substrates for the growth and proliferation of skin microbes, ultimately becoming integral components of their cellular structures (*59*). Microorganisms can acquire FFAs via two main strategies: *de novo* synthesis or uptake from the external environment. For instance, *C. acnes* are able to synthesize FFAs through the bacterial type II fatty acid synthesis (FAS II) pathway (*60*), while certain eukaryotes and actinomycetes rely on the type I fatty acid synthesis (FAS I) pathway (*61*). In contrast, auxotrophic species such as *Malassezia restricta* and *Corynebacterium jeikeium*, which lack the capacity for lipid biosynthesis, must rely on exogenous FFAs to support growth (*37, 62–64*). Notably, *de novo* fatty acid biosynthesis represents an energetically costly process, while environmental FFA uptake substantially reduces microbial metabolic demands (*59*).

Our analysis revealed that all microbial species coexisting with *C. acnes* were positively correlated with FFA levels (Figure 5a), and these species typically encoded genes involved in exogenous FFA uptake—such as fabH, the FakAB system, or acyl-CoA synthetases (FAA1/2/3/4) (Figure 5b). For example, Firmicutes such as *Staphylococcus spp.* possess the fatty acid kinase system (FakAB), which facilitates the recruitment and transport of exogenous FFAs across membranes (*65, 66*); *M. restricta* utilizes acyl-CoA synthetases (FAA1-4) to activate and incorporate external FFAs (*67*). The absence of fatty acid biosynthesis genes in *M. restricta* and *C. jeikeium* (Figure 5b) further supports their strict dependence on exogenous FFAs. In contrast, *C. acnes* and other *Cutibacterium* species retain a complete FAS II system, enabling autonomous FFA production.

**Figure 5.**
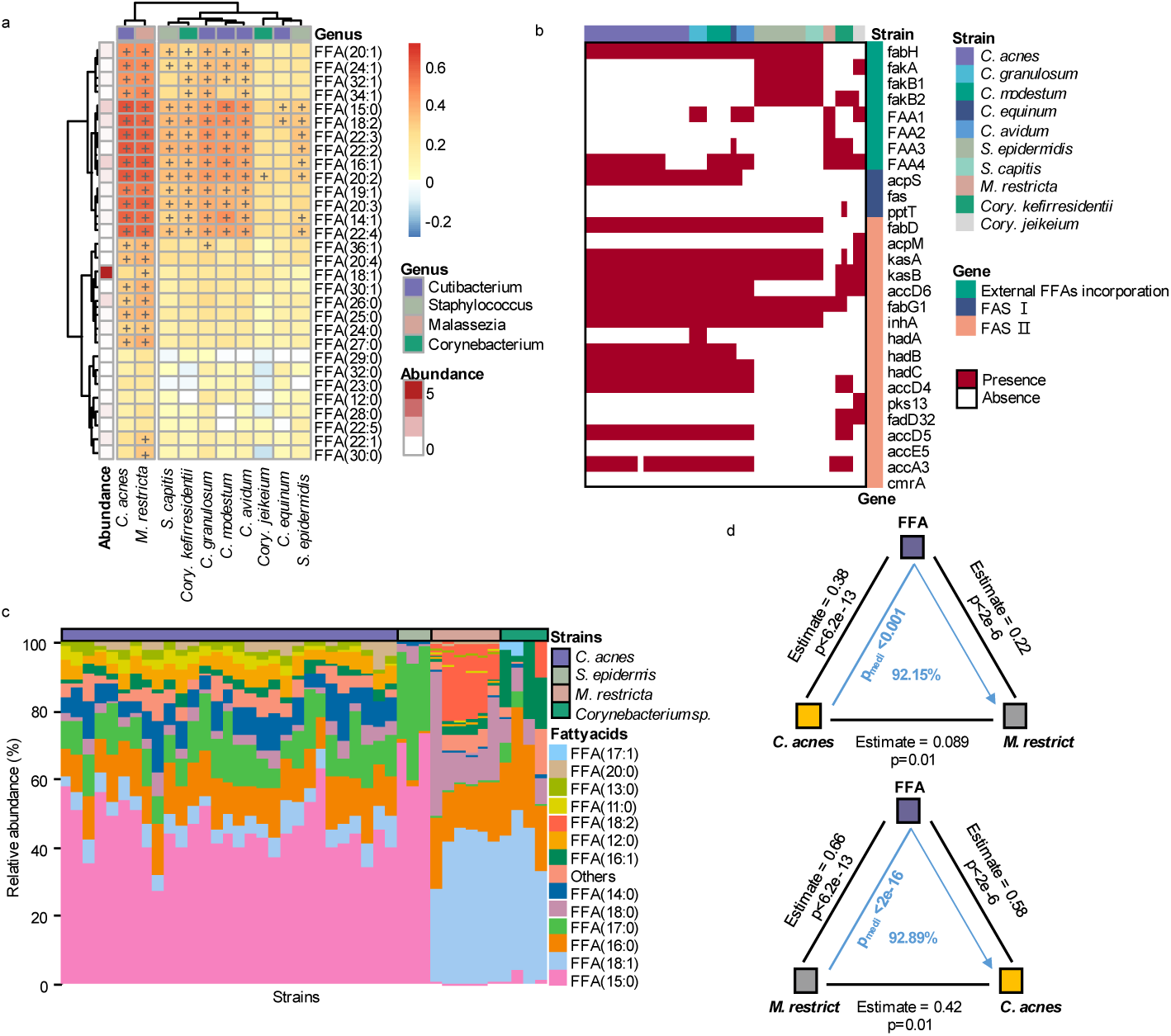
Free Fatty Acids as Central Regulators of the Skin Microecosystem. a Spearman correlations between FFAs and species. b Presence (red) or absence (white) of gene of free fatty acid transferase and synthase among strains. c The relative abundance (%) of free fatty acids in the strains. d *C. acnes*’s mediation effect on *M. restricta* via FFA. d *M. restricta*’s mediation effect on *C. acnes* via FFA.

Previous studies have identified C15:0 and C18:1 as the dominant FFAs incorporated into microbial membrane lipids (*67–71*), mirroring the most abundant FFAs detected on the skin surface (Figure 5c). Notably, FFAs derived from the host stratum corneum, predominantly long-chain saturated fatty acids (C16-C26), contribute minimally to the surface FFA pool (*72*), suggesting that the majority of surface FFAs originate from microbial metabolism of TGs rather than host-derived sources.

Mediation analysis revealed that *C. acnes* and *M. restricta* modulate each other’s abundance through FFA-mediated interactions, with FFAs acting as dominant intermediaries in this regulatory relationship. *C. acnes* exerted a significant indirect effect on *M. restricta* via FFAs (p = 0.008), while its direct effect was not significant (p = 0.736); FFA-mediated effects accounted for 92.1% of the total influence (Figure 6d). Similarly, *M. restricta* showed no significant direct effect on *C. acnes* (p = 0.75), but mediated 92.89% of its influence on *C. acnes* through FFAs (p < 2 × 10⁻¹⁶; Figure 6d). Taken together, these findings suggest that FFAs released from microbial TG metabolism, particularly by *C. acnes* and *M. restricta*, may function as “public goods” within the skin microecosystem. These are compounds or functions released into the extracellular environment that confer collective benefits to the microbial community (*73*). The defining characteristics of such public goods include shareability and communal impact: FFAs can be utilized both by their producers (e.g., *C. acnes*) and by auxotrophic microbes (e.g., *M. restricta*), and their cross-species flow supports the overall survival and stability of the skin microbiome.

**Figure 6.**
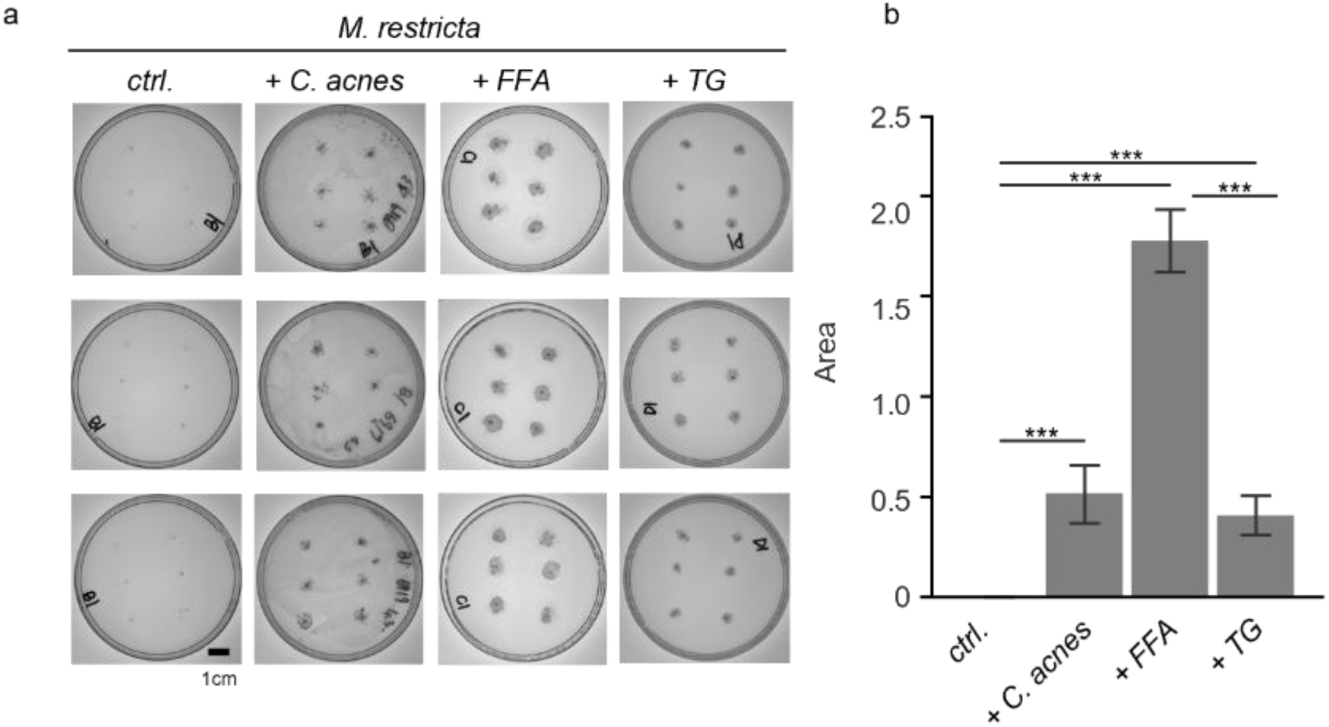
C. acnes create a metabolic niche for M. restricta. a *C. acnes* and *M. restricta* co-culture and lipid assimilation assay images. b Comparison of the growth areas of *M. restricta* in different culture mediums.

### *C. acnes* create a metabolic niche for *M. restricta*

To validate the metabolic interaction between the two lipophilic microbes, *C. acnes* and *M. restricta* were co-cultured *in vitro*. The results showed that *M. restricta* failed to grow in lipid-deficient basal medium but exhibited proliferation upon supplementation with either triglycerides (TG) or free fatty acids (FFA). Notably, growth was more robust in FFA-supplemented medium (Figure 6a and 6b), confirming our initial hypothesis that *M. restricta* preferentially utilizes FFAs over TGs.

Interestingly, although *M. restricta* failed to grow independently in lipid-deficient medium, it was able to grow successfully when co-cultured with *C. acnes* under the same conditions (Figure 6a and 6b). This finding reveals a metabolic interaction between *C. acnes* and *M. restricta*, suggesting that the presence of *C. acnes* may create a nutritional niche necessary for *M. restricta* survival. This exemplifies a niche construction mechanism driven by metabolic symbiosis within the skin microbiome.

## DISCUSSION

This study focuses on a population in a remote rural area of Yunnan Province, China.

By deciphering biotic compartments and the chemical landscape within an ecological theoretical framework, we revealed, at a high level of resolution, the role of environmental filtering, particularly that of the chemical microenvironment, in shaping the core skin microbiota. The core taxa and characteristic patterns (cutotype) of this non-industrialized population were highly consistent with those of industrialized groups in China (*9*). A recent study of the remote indigenous Yanomami community, a valuable population completely isolated from industrialized environments, showed that their skin microbiota exhibited much higher species diversity (*74*). Their most abundant bacterial taxa also include species of *Moraxella*, e.g., *M. osloensis*, previous identified linked to skin aging (*75*), was also present at high abundance in some individuals in our cohort. It is noteworthy that in this ancient population, *Malassezia restricta* was correlate inversely with *Moraxella* and positively with *Cutibacterium* (*74*), fully consistent with our data-driven clustering and niche differentiation analysis, placed *Malassezia restricta* and *Cutibacterium* along the same metabolic panel, but the opposite to *Moraxella* panel (Figure 3a). Furthermore, the relative abundance of *M. restricta* was negatively correlated with overall bacterial richness and diversity, which also aligns with our observations. These findings reveal stable and conserved skin microbial interactions across different populations, indicating fundamental ecological machinery in this ecosystem.

The host skin functions as a “soil” that selects its colonists, with lipids rather than water-soluble compounds, serving as the key environmental filter. This lipid-driven filtering profoundly shapes the distribution of keystone species and the modular structure of the microbial community. *Cutibacterium acnes*, with its broad lipid metabolic capacity (including diverse lipases), holds a definitive competitive advantage that underpins its keystone status. In high-lipid environments (e.g., during adolescence), the competitive dominance of *C. acnes* suppresses community diversity (*32, 33*). As sebum levels decline, the competitive pressure exerted by *C. acnes* diminishes, releasing ecological niches and allowing diversity to increase, consistent with age-dependent diversity elevation (*8*). Notably, under declining lipid conditions, we observed a transient increase in *Staphylococcus epidermidis*, a moderately lipophilic and fast-growing species (Fig. S4a). However, once lipid availability falls below a critical threshold, *S. epidermidis* also declines. In contrast, *Moraxella osloensis*, a non-lipophilic, oligotrophic, and rapidly proliferating organism (*39–42*), gains a competitive advantage under low-lipid conditions, leading to its dominance at the opposite end of the lipid spectrum. These two keystone species occupy largely non-overlapping ecological niches, demonstrating clear niche partitioning that minimizes direct competition and results in dual abundance peaks across the lipid gradient. This ecological specialization provides a mechanistic explanation for previously proposed “cutotypes” derived from data-driven approaches, which have since been validated across multiple independent cohorts (*10, 11*).

Another intriguing finding is that microbial community diversity markedly declines when either *C. acnes* or *M. osloensis* achieves absolute dominance. In contrast, a transitional state of relative equilibrium between the two is associated with increased diversity. This pattern aligns with the core prediction of the Intermediate Disturbance Hypothesis (IDH). The intermediate “dual-species equilibrium” represents a state of moderate disturbance that mitigates competitive exclusion, creates new ecological opportunities, and facilitates multispecies coexistence, a phenomenon analogous to grassland ecosystems where grazing suppresses dominant plants and enhances biodiversity (*76*). This insight may extend to common dysbiosis-related disorders. In acne vulgaris, the clear dominance of *C. acnes* reduces microbial diversity and suppresses other taxa (*77*). Similarly, at the other extreme, the rapid proliferation of *M. osloensis* under low-lipid conditions, linked to skin aging (*75*), parallels the behavior of “weedy” species such as *Clostridioides difficile* in the gut, which rapidly colonize when niche vacancies appear (*78*). Therefore, leveraging IDH theory to induce a state of “intermediate disturbance” and promote transitional diversity peaks may offer a novel direction for therapeutic strategies.

The application of ecological theory to understand the human microbiome is not new. As early as 2012, a systematic review explored how metacommunity theory explains microbiome variation, proposing that community assembly is governed by dispersal, diversification, environmental selection, and ecological drift (*79*), and concepts such as “everything is everywhere, but the environment selects” (*80*) have shaped our understanding of microbial assembly. Nevertheless, practical efforts to maintain and restore microbial communities are often hindered by a lack of operable design principles (*81*). To address this, a general framework for managing ecosystem services has been proposed (*82*), aiming to systematically advance ecology-informed health interventions. Within this framework, our study has made several key advances:

First, we identified key ecosystem service providers (ESPs) and elucidated their functional mechanisms. Through lipid-microbe correlation and mediation analyses, *Cutibacterium acnes* and *Malassezia restricta* were identified as core functional species. Utilizing their extensive lipase arsenal (notably triglyceride hydrolase activity), they dominate the core metabolic process on the skin surface: the conversion of triglycerides (TGs) into free fatty acids (FFAs). These species collectively maintain a dynamic balance in the TG–FFA ratio (accounting for ∼85% of total skin lipids), while other lipase-producing species (e.g., *Staphylococcus epidermidis*) contribute minimally. As key “public goods” in the microbial ecosystem, FFAs support the growth of numerous lipid-auxotrophic microorganisms (e.g., *M. restricta*), thereby sustaining broad microbial interactions. *C. acnes* and *M. restricta* regulate each other’s abundance via FFAs, forming a tight symbiotic association. Importantly, *in vitro* coculture experiments confirmed that *C. acnes* establish a survival niche for *M. restricta*, significantly promoting the latter’s growth in environments where it cannot thrive independently, demonstrating unidirectional metabolic cross-feeding and providing causal validation for this nutrient niche construction.

Second, our study revealed how community context influences ESP function. *C. acnes* dominate in high-lipid environments (e.g., sebum-rich regions), maximizing its ecosystem service output (FFA production). Concurrently, skin lipids exert selective pressure on *C. acnes*, which indirectly influences other taxa through cross-feeding and other interactions. This mechanism, analogous to short-chain fatty acid (SCFA)-mediated interactions in the gut(*83, 84*), operates in a highly specialized context: the lipid-rich, acidic, and dry skin environment contrasts sharply with the carbohydrate-rich, neutral-pH gut (*36*) or the mucus-dominated lung (*85*), selecting for highly adapted taxa such as *C. acnes*, which thrives in anaerobic, lipid-rich follicles. We also uncovered a “resource diversity paradox”: increased lipid diversity correlates with reduced microbial diversity, indicating that lipids act as selective filters rather than general nutrients, opposite to the gut, where resource diversity typically enhances microbial diversity. These findings underscore the importance of organ-specific ecological principles in microbiome assembly and function. Although the spatiotemporal dynamics of ESP operation require further investigation, our study has made substantial progress in identifying ESPs, deciphering functional mechanisms, and assessing environmental regulation, laying a foundation for ecology-informed interventions targeting the skin microbiome.

In summary, this study establishes an ecological framework for skin microbiome assembly. Our findings not only extend the application of ecological theory to human microbial systems but also provide important insights into the pathogenesis of skin disorders and the development of novel therapies. Future studies should integrate longitudinal tracking and functional experiments to further elucidate the molecular mechanisms underlying keystone species dynamics and to design individualized strategies accounting for ethnic variations. These efforts will advance skin microbiome research from a descriptive science toward an interventional one, ultimately enabling ecology-based precision management of skin health.

## MATERIALS AND METHODS

### Study population and sample collection

170 female healthy volunteers, who were 20 to 60 years old, were recruited from the general population in Yunnan in November in 2023. Medical and medication history was obtained for each individual by questionnaires. Subjects with any history of skin diseases and intake of systemic or local antibiotics in the past 6 months were excluded. To maximize microbial and metabolites skin load, each subject was instructed to wash the face only with tap water and to refrain from the application of any skin-care or cosmetic products on the sampling day before sampling. The cheek site was sampled for each subject. Study personnel wore sterile gloves for each sample collection. Samples were collected in a temperature and humidity-controlled room at 20 °C and 50% humidity.

The samples for microbiome analysis were collected by using sterile cotton-tipped dry swabs that had been pre-moistened with GDS-01A solution (GeWei Bio-Tech (shanghai) Co. Ltd, Shanghai, China). Swabs were rubbed firmly on the cheek for 40 times to cover a facial surface area of 4 cm2. Then, the swab head was fractured, placed in a sterilized 1.5 mL centrifuge tube, and stored at − 80 °C.

The samples for skin surface metabolome and lipidome analysis were collection by using DS tape DS tape (22mm, CuDerm, Dallas, TX, USA). The procedure for collecting skin metabolites with DS tape was adapted from methods previously described (*86, 87*) with some modifications. The tape was applied to designated cheek region using sterile forceps, pressed and stripped at a defined frequency using a constant pressure rod. After stripping, the tape was removed with forceps, placed in a 2-mL sterile centrifuge tube (adhesive side in), and immediately stored at −80℃.

### DNA extraction and metagenomic sequencing

Total DNA from a skin swab was isolated using VAMNE Magnetic Pathogen DNA Kit (Vazyme, Nanjing, PRC) according to the manufacturer’s protocols. The purity and concentration of the extracted total DNA were estimated by NanoDrop™ One and Qubit Flex Fluorometer (Thermofisher Scientific, Waltham, MA, USA). The extracted DNA was amplified by using Discover-sc Single Cell WGA Kit (Vazyme, Nanjing, PRC), then the multiple displacement amplification (MDA) product was quantified by Qubit Flex Fluorometer (Thermofisher Scientific, Waltham, MA, USA). DNA-seq library was prepared with approximately 200ng of MDA product using VAHTSTM Universal DNA Library Prep Kit for Illumina® (Vazyme, Nanjing, PRC). Briefly, the DNA was fragmented and end-repaired. Then the A tailing and adapter ligation were performed with the end-repaired DNA. Finally, the purified, adapter-ligated DNA was amplified. The library quality and concentration were assessed by utilizing a D1000 Screen Tape on an Agilent 4200 Tapestation. Accurate quantification for sequencing applications was determined using the qPCR-based KAPA Biosystems Library Quantification kit (Kapa Biosystems, Inc., Woburn, MA). Each library was diluted and pooled equimolar prior to circularization. Paired-end (PE150) sequencing was performed on MGI DNBSEQ-T7 platform.

### Sequencing and quality control

We processed and sequenced 170 samples and four negative controls (i.e., new sterile swabs). Sequence adapters were removed from the raw reads using AdapterRemoval (v2.3.1) (*88*) and quality filtering and human DNA read removal was performed using Kneaddata (v0.7.4) as previously described (*22*). Sequencing quality information for the samples is shown in Supplementary Table 6. Following quality filtering, an average of 7,612,776,128 paired-end reads were retained per sample. Following taxonomic classification of the reads using Kraken2 (v2.0.7-beta; kmer length = 35, confidence score threshold = 0) (*89*), then species-level abundance estimation using Bracken (v2.6.1; threshold for filter = 0) (*90*), putative contaminants were identified with the R package “decontam” based on the prevalence mode with 0.1 as the significance threshold (*91*). This method deemed 62 species to be contaminants (Supplementary Table 6) and species that could be mapped to the contaminant species of each sample (Supplementary Table 6) were removed.

### Sloan neutral model prediction

To assess whether the microbial communities exhibit either a neutral or a niche-based community assembly process, the Sloan neutral model analysis (*92*) was performed for microbiome as described previously for the skin microbiota (*23*) to predict the relationship between OTU detection frequency and their relative abundance across the wider metacommunity (*92*). In this model, the parameter R2 represents the overall fit to the neutral model (*92*). Calculation of 95% confidence intervals around all fitting statistics was done by bootstrapping with 1000 bootstrap replicates. The Sloan model prediction and statistics were performed in R (*23*).

### Prediction of relative abundance of *C. acnes*, *M. restricta* and *S. epidermidis* from skin surface metabolomic data

The 170 healthy individuals were randomly divided into training and validation sets. In the training set, the ElasticNet regression model was applied to log-transformed data of metabolomic data using the glmnet (*93*) R package (version 4.1-8). In *C. acnes*, *M. restricta* and *S. epidermidis* measurement data, only individuals with blank values less than 10 were kept, and features that were not detected in all the kept individuals were further excluded. The model was built with alpha values of 0.1–0.9. And the lambda values were selected using a 3-fold cross-validation on the training sets. The models with the smallest mean absolute error (MAE) in the validation set were selected as the final models (*94*).

### Extraction of hydrophilic and hydrophobic metabolites

To extract hydrophilic compounds, each sample was thawed on ice. The tape was then transferred to a corresponding labeled centrifuge tube, followed by the addition of 3 mL of 70% methanol containing internal standards. The mixture was vortexed at 2500 r/min for 5 min, and the tape was removed. After resting on ice for 15 min, the sample was centrifuged at 3000 r/min at 4 °C for 10 min. Then, 1.2 mL of the supernatant was transferred to a new labeled tube and left at −20 °C for 30 min. Subsequently, the sample was centrifuged again at 12000 r/min at 4 °C for 3 min. Finally, 120 μL of the supernatant was transferred to an LC vial with an insert for LC-MS/MS analysis.

To extract hydrophobic compounds, each sample was thawed on ice and mixed with 1 mL of lipid extraction solvent containing internal standards (methyl tert-butyl ether: methanol = 3:1, v/v). The mixture was vortexed for 15 min, followed by the addition of 200 μL of water and vortexed for 1 min. Then, the sample was centrifuged at 12,000 r/min at 4 °C for 10 min. A total of 500 μL of the upper phase was transferred to a new labeled centrifuge tube and evaporated to dryness. The residue was re-dissolved in 200 μL of lipid reconstitution solvent (acetonitrile: isopropanol = 1:1, v/v), vortexed for 3 min, and centrifuged at 12,000 r/min for 3 min. The supernatant was collected and subjected to LC-MS/MS analysis.

### LC analysis of hydrophilic and hydrophobic compounds

The sample extracts of hydrophilic compounds were analyzed using the ultra-performance liquid chromatography (UPLC) of a LC-MS/MS system (ExionLC AD, https://sciex.com.cn/, MS, QTRAP® 6500+ System). The samples were injected onto a Waters ACQUITY UPLC HSS T3 C18 (1.8 µm, 2.1 mm × 100 mm). The column temperature, flow rate and injection volume were set 40 °C, 0.4 mL/min and 2 μL, respectively. Mobile phase: Phase A is ultrapure water (0.1% formic acid), and Phase B is acetonitrile (0.1% formic acid). Elution gradient: At 0 min, water/acetonitrile (95:5 V/V); at 2.0 min, 80:20 V/V; at 5.0 min, 40:60 V/V; at 6.0 min, 1:99 V/V; at 7.5 min, 1:99 V/V; at 7.6 min, 95:5 V/V; at 10.0 min, 95:5 V/V.

The sample extracts of hydrophobic compounds were also analyzed using the same UPLC (ExionLC™ AD, https://sciex.com.cn/, MS, QTRAP® 6500+ System). The samples were injected onto a Thermo Accucore™ C30 column (2.6 μm, 2.1 mm × 100 mm). The column temperature, flow rate and injection volume were set 45 °C, 0.35 mL/min and 2 μL, respectively. Mobile phase: Phase A, acetonitrile/water (60/40, V/V) (containing 0.1% formic acid, 10 mmol/L ammonium formate); Phase B, acetonitrile/isopropanol (10/90, V/V) (containing 0.1% formic acid, 10 mmol/L ammonium formate).Gradient elution program: At 0 min, A/B = 80:20 (V/V); at 2 min, 70:30 (V/V); at 4 min, 40:60 (V/V); at 9 min, 15:85 (V/V); at 14 min, 10:90 (V/V); at 15.5 min, 5:95 (V/V); at 17.3 min, 5:95 (V/V); at 17.5 min, 80:20 (V/V); at 20 min, 80:20 (V/V).

### MS/MS-based analysis of hydrophilic and hydrophobic compounds

Triple quadrupole (QQQ) scans with linear ion trap (LIT) were acquired using the LC-MS/MS system (Shim-pack UFLC SHIMADZU CBM A system, MS, QTRAP® 6500+ System). This system was equipped with an electrospray ionization (ESI) Turbo Ion-Spray interface, which could be operated in positive and negative ion modes and controlled by Analyst 1.6.3 software package (Sciex).

The parameters of the ESI source operation for hydrophilic compounds were set as: ion source, turbo spray; source temperature, 500 °C; ion spray voltage, 5500 V in the positive ion mode or −4500 V in the negative ion mode; collision-activated dissociation, high; ion source gas I, 55 psi; ion source gas II, 60 psi; curtain gas, 25 psi.

The quantification of metabolites was accomplished using targeted multiple reaction monitoring (MRM) approach.

The parameters of the ESI source operation for hydrophobic compounds were set as: ion source, turbo spray; source temperature, 500 °C; ion spray voltage, 5500 V in the positive ion mode or −4500 V in the negative ion mode; collision-activated dissociation, medium; ion source gas I, 45 psi; ion source gas II, 55 psi; curtain gas, 35 psi.

The quantification of metabolites was accomplished using targeted multiple reaction monitoring (MRM) approach.

### Data analysis of skin hydrophilic and hydrophobic metabolites

The MS/MS data were processed by Analyst 1.6.3 software package (Sciex). The reproducibility of metabolite extraction and detection were judged by total ion current and multiple peaks of MRM quantifications. According to the retention time and mass-to-charge ratio, the identification of both hydrophilic and hydrophobic metabolites was performed using a home-made metadata database and other existing metabolomic databases, including HMDB 4.0 (https://www.kegg.jp/<u>), Mona</u> (https://mona.fiehnlab.ucdavis.edu/<u>), MassBank</u> (http://www.massbank.jp/).

For quantification of metabolites, the metabolomic data was analyzed using MultiQuant software package 3.0.2 (Sciex), which automatically integrated and calibrated the chromatographic peaks. The IBA of each metabolite was calculated by the peak area of each chromatographic peak.

### Lipid assimilation assays

*M. restricta* cells grown for 7 days on ready-to-use Pityrosporum agar plates were harvested and transferred into sterile 5 mL centrifuge tubes. The cells were washed three times with sterile phosphate-buffered saline (PBS) by centrifugation and subsequently resuspended in 3 mL PBS to obtain a uniform cell suspension.

Three types of culture media were prepared:

- Medium B: Sabouraud broth supplemented with 3% Sea Plaque GTG (low-melt) agarose;
- Medium C: Medium B supplemented with 2 mL of oleic acid;
- Medium D: Medium B supplemented with 2 mL of triolein.

Each medium was melted, equilibrated to 45°C, and poured into 90 mm Petri dishes. After solidification, 5 µl of the bacterial suspension was added to the plates, spotting 6 times. Once the spots dried, plates were wrapped in gas-permeable parafilm and incubated in the Three-phase incubator set at 2% O2, 5% CO2, and 34°C for 7 days.

### Co-culture experiments

*M. restricta* cultured for 7 days on ready-to-use Pityrosporum agar plates and *C. acnes* cultured for 4 days on ready-to-use Columbia blood agar plates were harvested and transferred into sterile 5 mL centrifuge tubes. Each strain was washed three times with sterile PBS by centrifugation, then resuspended in 3 mL sterile PBS to obtain separate monoculture suspensions. Under anaerobic conditions, 100 μL of the *C. acnes* suspension was spread evenly onto the surface of the agar plates and air-dried. Then, 5 μL of the *M. restricta* suspension was spotted onto the same plate using a 10 μL pipette, with six replicate spots per plate. After the spots had dried, the plates were sealed with gas-permeable parafilm and incubated in a tri-gas incubator at 32°C under conditions of 2% O₂ and 5% CO₂ for 7 days.

## ACKNOWLEDGMENT

We sincerely thank Yatsen Global Innovation R&D Center and EveLab Insight (a subsidiary of Meitu Inc.) for their valuable support. Their contributions have greatly enhanced our research efforts, and we are truly grateful for their trust and collaboration.

## AUTHOR CONTRIBUTIONS

Huizhen Chen contributed to investigation, formal analysis, visualization, experimental validation, and drafted the original manuscript. Wei Luo assisted with cohort establishment, biospecimen collection, project administration and investigation. Zhiming Guo & Zhiming Li provided scientific discussion and participated in manuscript preparation. Jianju Feng, Shensong Cao, and Qi Wang assisted with biospecimen collection. Zhenglong Gu contributed to scientific discussion and recommendations. Jean Krutmann and Jiucun Wang provided supervision and critical review of the manuscript. Lianyi Han contributed scientific supervision, resources, and manuscript review. Yangfan Guo provided supervision, resources, funding acquisition, project administration, and manuscript review. Jingjing Xia oversaw the study through conceptualization, supervision, secured funding, managed project administration, and participated in drafting the original manuscript. All authors read, edited and approved the paper.

---

**Figure S1.**
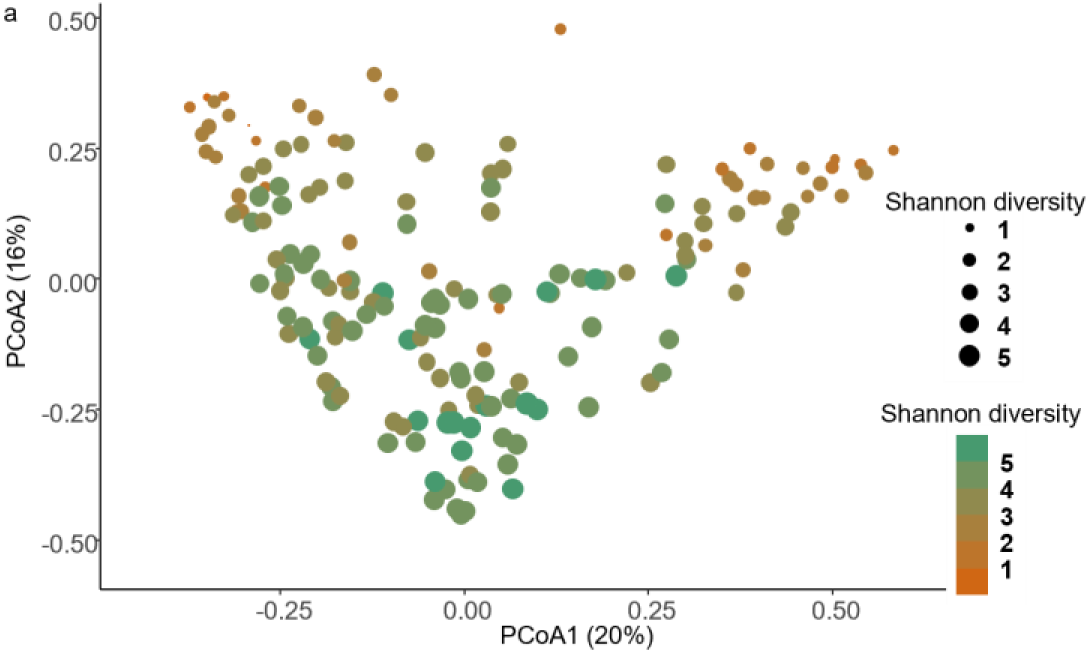
a Projection of 170 samples using PCoA analysis (Bray-Curtis dissimilarity).

**Figure S2.**
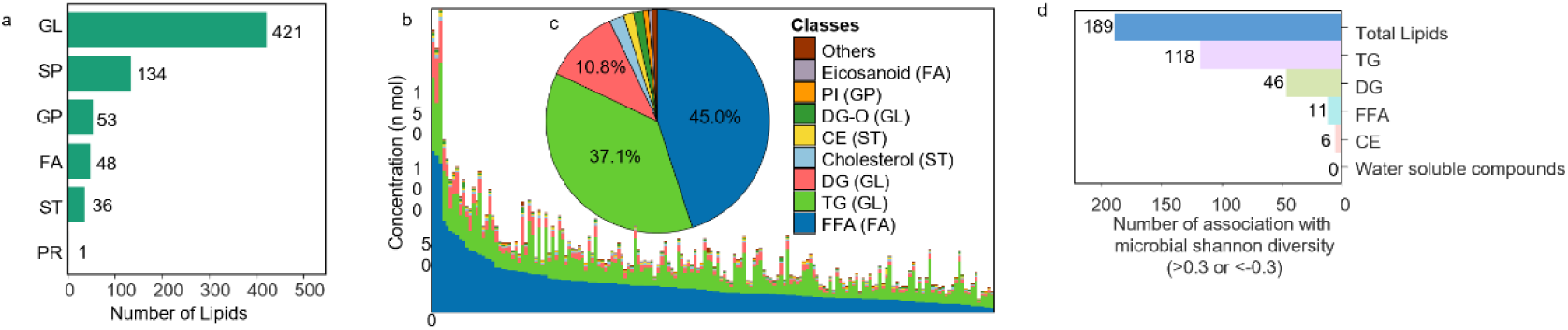
a The bar plot represents the counts of each class across all samples. b The compositions of the top 8 lipid classes, ranked by their average concentration across all samples. c Proportion of top 8 lipid classes. d Distribution of association counts (absolute Spearman correlation > 0.3) between metabolite species and PCoA1 coordinates in Figure S1a.

**Figure S3.**
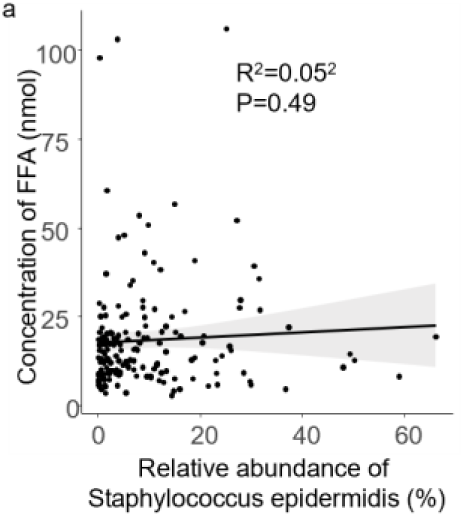
a Correlation between relative abundance of *Staphylococcus epidermidis* and concentration of FFA in the population.

