## SupplementaryFigues for "Integrative Multi-omics Deciphers the Assembly Rules of the Human Skin Microbiome and Highlights Lipid-Mediated Niche Partitioning"

### Supplementary Figures

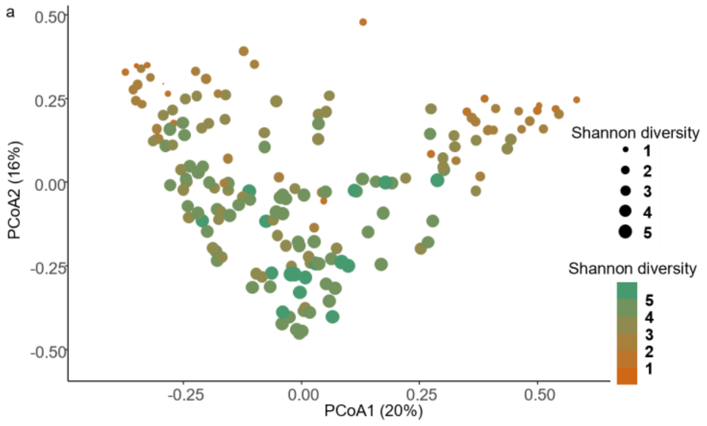


Figure S1. a Projection of 170 samples using PCoA analysis (Bray-Curtis dissimilarity).

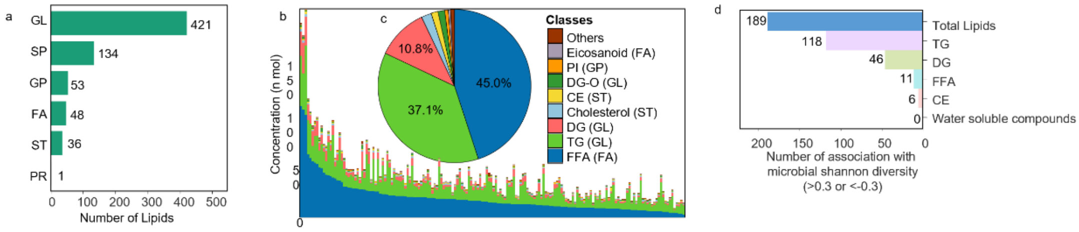


Figure S2. a The bar plot represents the counts of each class across all samples. b The compositions of the top 8 lipid classes, ranked by their average concentration across all samples. c Proportion of top 8 lipid classes. d Distribution of association counts (absolute Spearman correlation > 0.3) between metabolite species and PCoA1 coordinates in Figure S1a.

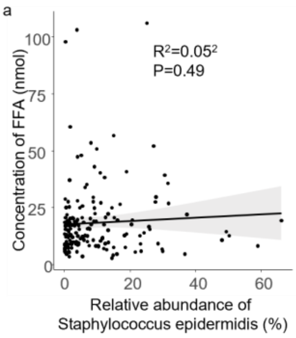


Figure S3. a Correlation between relative abundance of *Staphylococcus epidermidis* and concentration of FFA in the population.
